# AXL-GAS6/PROS1 Interaction: A Critical Switch Between Aberrant- and Healthy Repair Following Alveolar Lung Injury

**DOI:** 10.1101/2025.04.07.647567

**Authors:** Devona Soetopo, Christoph H. Mayr, Katrin Fundel-Clemens, Fidel Ramirez, Coralie Viollet, Alec Dick, Werner Rust, Diana Santacruz, Yvette Hoevels, Christopher J. Applebee, Sam Legg, Anastasia Funk, Gisela Schnapp, Julian Padget, Benjamin Strobel, Matthew J. Thomas, Stephen G. Ward, Banafshé Larijani, Kerstin Geillinger-Kästle

**Affiliations:** Immunology and Respiratory Diseases Research, Boehringer Ingelheim Pharma GmbH & Co. KG, Biberach an der Riss, Germany; Computational Biology and Digital Sciences, Boehringer Ingelheim Pharma GmbH & Co. KG, Biberach an der Riss, Germany; Medicinal Chemistry, Boehringer Ingelheim Pharma GmbH & Co. KG, Biberach an der Riss, Germany; Drug Discovery Sciences, Boehringer Ingelheim Pharma GmbH & Co. KG, Biberach an der Riss, Germany; Consortium for Precision Health and Department of Life Sciences, University of Bath, Bath, United Kingdom; Department of Computer Science, University of Bath, Bath, United Kingdom

**Keywords:** Aberrant alveolar repair, Epithelial injury, Pulmonary fibrosis, AAV-DTR, AXL-GAS6-PROS1

## Abstract

**Rationale:** Idiopathic pulmonary fibrosis (IPF) is a progressive lung disease characterized by aberrant alveolar repair and excessive fibrosis. The TAM-family receptor tyrosine kinase AXL, activated by GAS6 and PROS1, is implicated in tissue remodeling, but ligand-specific AXL signaling during alveolar repair remains poorly defined.

**Objectives:** To investigate ligand specific AXL signaling in IPF and how it impacts epithelial proliferation and repair after alveolar injury *in-vivo* and *in-vitro*.

**Methods:** Single cell RNA sequencing was utilized to understand cell specific expression patterns in IPF patients, followed by functional studies in primary human cell culture and functional spatial digital profiling (FuncOmap) analysis in patient tissue. Longitudinal assessment of repair process after alveolar-specific injury *in-vivo* was used to complement the *in-vitro* approach.

**Results:** AXL expression showed enrichment in basal and aberrant basaloid cells of IPF patients. *In-vitro* GAS6 increased proliferation of basal cells, while PROS1 counteracted this effect. FuncOmap analysis demonstrates direct *in-situ* interactions between AXL and both ligands, providing evidence for biological relevance. Investigating longitudinal repair processes *in-vivo* revealed dynamic regulation of AXL ligands as well as AXL.

**Conclusions:** These findings highlight the importance of ligand-specific AXL signaling in lung repair and suggest that it dysregulation may contribute to IPF pathogenesis, offering potential therapeutic targets for restoring normal repair processes.

## INTRODUCTION

Idiopathic pulmonary fibrosis (IPF) is a progressive lung disease characterized by aberrant epithelial repair leading to irreversible fibrosis, predominantly affecting older individuals. Despite extensive research, the underlying mechanisms driving IPF remain poorly understood, and current treatments only slow down disease progression^1^. While the precise etiology of IPF remains unclear, it is hypothesized to result from repeated alveolar epithelial injury followed by aberrant repair processes that culminate in fibrosis^2–4^. Understanding the molecular and cellular mechanisms that regulate normal alveolar repair is critical for identifying therapeutic targets to restore homeostasis and prevent fibrosis.

Recent advances in single-cell RNA sequencing (scRNA-seq) have revealed the cellular heterogeneity of IPF lungs, identifying basal and aberrant basaloid cells enriched in IPF and implicated in pathological remodeling^5–7^. However, the signaling pathways that govern epithelial cell behavior during repair and their dysregulation in IPF remain to be fully elucidated.

AXL, a receptor tyrosine kinase of the TAM family (TYRO3, AXL, MERTK), regulates cellular proliferation, tissue repair, and immune modulation. Activated by its ligands GAS6 and PROS1, AXL initiates downstream signaling cascades, including the PI3K/AKT and MAPK pathways, to promote cell survival and proliferation. While GAS6 exhibits the highest affinity for AXL, PROS1 predominantly binds to TYRO3 and MERTK^8,9^. Although the interaction between PROS1 and AXL has been debated^10–15^, our findings demonstrate this interaction and suggests its potential contribution to lung repair process. While AXL signaling has been implicated in tissue repair and fibrotic diseases^16–20^, its role in IPF and its contribution to epithelial cell dysfunction remain underexplored. Gaining a deeper understanding of the roles of AXL and its ligands in IPF pathogenesis could unlock new avenues for therapeutic intervention.

Here, we integrate scRNA-seq data from IPF lungs with functional studies in primary human small airway epithelial cells and a mouse epithelial injury model to define the roles of AXL, GAS6, and PROS1 in lung repair. We show that AXL and its ligands are upregulated in IPF lungs and exert ligand-specific effects on epithelial proliferation and repair. These findings identify dysregulated AXL signaling as a potential driver of impaired epithelial repair in IPF.

## RESULTS

### AXL and its ligands GAS6 and PROS1 are upregulated in IPF patients

Using an integrated scRNA-seq of human IPF atlas^5^, we found that *AXL* and its ligands *GAS6* and *PROS1* were upregulated in the epithelial cells of IPF patients. Higher expression of *AXL* was observed particularly in basal cells and aberrant basaloid cells, a cell type that is found predominantly in IPF^5–7^. The ligands *GAS6* and *PROS1* were highly expressed in AT-1 cells and the aberrant basaloid cells (**Figure 1A**). Consistently, elevated levels of GAS6 and PROS1 were also measured in the bronchoalveolar lavage fluid (BALF) of IPF patients compared to non-diseased controls and chronic obstructive pulmonary disease (COPD) patients, with PROS1 being 20-30-fold higher than GAS6 (**Figure 1B**).

**Figure 1.**
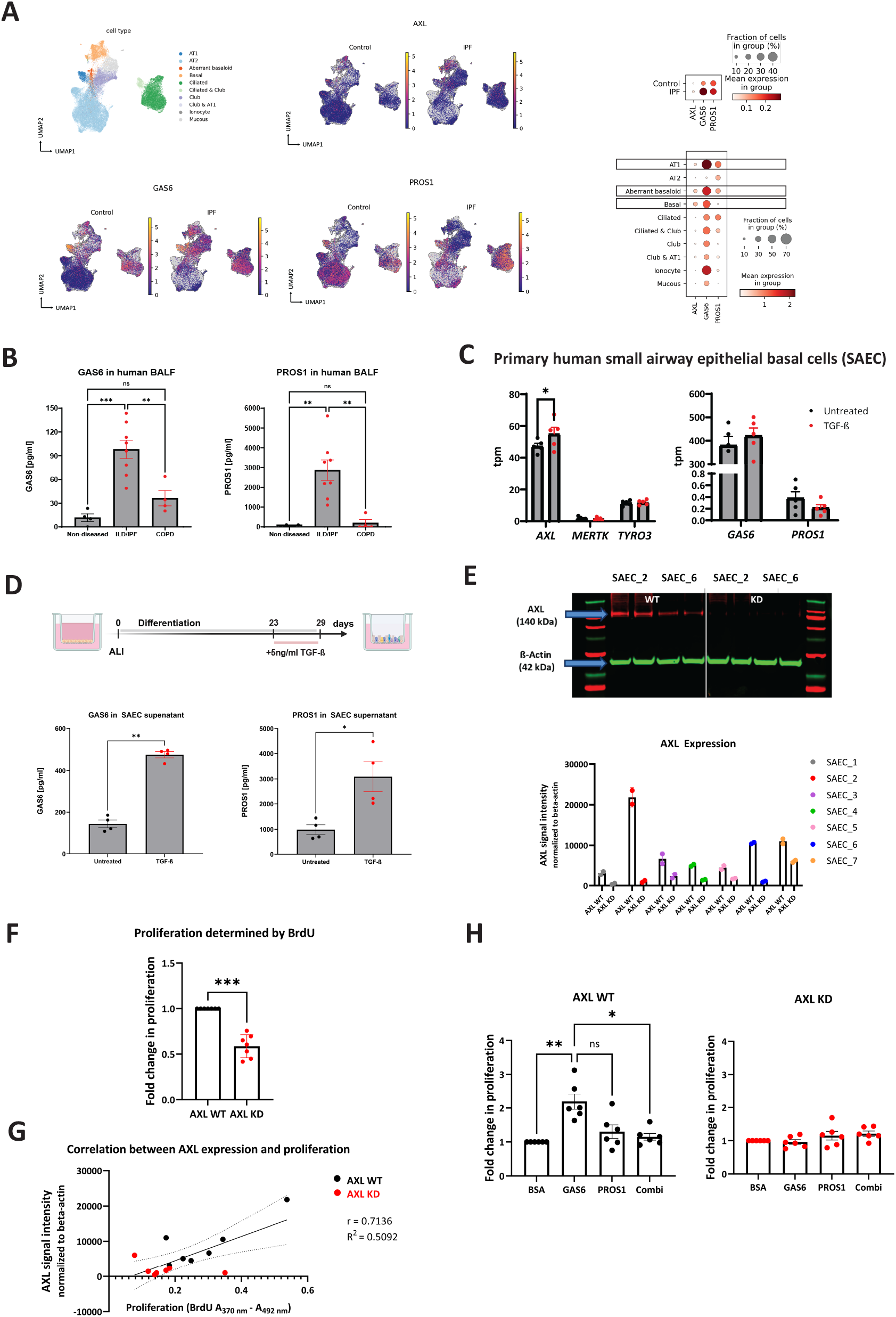
*AXL* is predominantly expressed in basal and aberrant basaloid cells, promoting proliferation via AXL-GAS6 signaling. **(A)** UMAP plots visualize *AXL* and its ligands, *GAS6* and *PROS1* normalized expression in the epithelial cell compartment of an integrated scRNA-seq IPF atlas. Upper dot plot shows quantification of *AXL* and its ligands expression in epithelial cells of IPF patients and the non-diseased controls. Lower dot plot presents the combined expression of AXL and its ligands from IPF and non-diseased controls across various epithelial cell types in the lung. **(B)** GAS6 and PROS1 concentration in human BALF of non-diseased control, ILD/IPF-, and COPD patients. **(C)** TAM receptors and their ligands expression in SAEC under submerged culture condition with and without 5 ng/ml TGF- ß treatment. **(D)** SAEC were differentiated for 23 days. On day 23 post ALI, 5 ng/ml rhTGF-ß was added to the culture for 6 days. On day 29 post ALI, GAS6 and PROS1 concentration in the SAEC cell culture supernatant was measured. **(E)** Representative western blot image validating knockout of AXL via CRISPR/Cas9 in 2 different donors. AXL (140 KDa, red bands), loading control ß-Actin (42 KDa, green bands), and M is protein marker. The graph shows quantification of AXL expressions of each donor before and after CRISPR/Cas9 mediated knockdown. Experiments were done in duplicate. **(F)** Proliferation of AXL WT and AXL KD SAEC as assessed by BrdU in all donors 48 hours post seeding. **(G)** Correlation between AXL expression in (F) and SAEC proliferation in (G). **(H)** Proliferation of AXL WT and AXL KD SAEC as assessed by BrdU 48 hours post treatment with rhGAS6/ rhPROS1/ combination of both proteins. WT: Wildtype; KD: Knockdown. Data are shown as mean ± SEM of n= 4 – 8. P >0.05 (ns/ non-significant); P ≤ 0.05 (*); P ≤ 0.001 (***).

Since *AXL* was predominantly expressed in basal cells of IPF patients, we sought to determine whether *AXL* was also expressed in primary human small airway epithelial basal cells (SAEC), which provide a tractable human basal epithelial model for mechanistic studies. While aberrant basaloid cells represent a disease-specific population in IPF, their limited accessibility currently precludes their use in functional *in-vitro* assays.

mRNA-seq of SAEC unveiled significant expression of *AXL* under submerged culture conditions, whereas expression of the other TAM receptors, *MERTK* and *TYRO3*, was low. Additionally, the ligand *GAS6* appeared to be highly expressed, whereas *PROS1* demonstrated minimal expression. Treatment with TGF-ß, a key profibrotic cytokine in IPF^21^, slightly enhanced *AXL* and *GAS6* expression, while *PROS1* decreased (**Figure 1C**). Under air-liquid interface (ALI) culture, TGF-ß treatment led to increase in the secretion of GAS6 and PROS1, which reflected the levels of both proteins in the BALF of IPF patients (**Figure 1D**).

Since basal cell proliferation is a critical early component of epithelial repair following injury, we used proliferation as a functional readout of epithelial repair capacity downstream of AXL signaling. To assess the role of AXL in this process, AXL knockout (KO) was performed in SAEC using CRISPR/Cas9. Baseline AXL expression varied across seven donors, with Donor SAEC_2 having the highest AXL expression and Donor SAEC_1 the lowest. Additionally, post-KO quantification of AXL expression showed that the KO efficiency differed across donors, thereby achieving a bulk knockdown (KD). To account for this variation at the start of the experiment, we assessed the AXL expression (**Figure 1E**).

Functionally, AXL-deficient SAEC displayed a significant reduction in proliferation compared to wild-type (WT) cells (**Figure 1F**). Moreover, the differential AXL expression among SAEC donors positively correlated with their proliferation capacities (**Figure 1G**), supporting a direct role of AXL signaling in basal epithelial proliferation. Consistent with this, stimulation of SAEC with AXL ligand rhGAS6 (400 ng/ml) significantly enhanced their proliferation capacity, whereas rhPROS1 (1200 ng/ml) alone had no impact on proliferation. However, combination treatment with 400 ng/ml rhGAS6 and 1200 ng/ml rhPROS1 abolished pro-proliferative effect of rhGAS6, suggesting rhPROS1 competing with rhGAS6 for AXL binding (**Figure 1H**). Conversely, none of these treatments affected proliferation in AXL KD SAECs, confirming that the observed effects were AXL-dependent (**Figure 1H**). The ligand concentrations used reflect those measured in IPF BALF, where PROS1 levels substantially exceed GAS6 (**Figure 1B**) and were further supported by dose response curve analysis (**Figure S1**).

### AXL-GAS6/ PROS1 interactions determined by Surface Plasmon Resonance and functional digital spatial profiling in IPF patient samples

Although AXL-GAS6 interactions are well described, PROS1 is generally considered a ligand for MERTK and TYRO3^8,9^. Given our data suggesting competitive effects of GAS6 and PROS1 on AXL signaling (**Figure 1H)**, we investigated the direct interaction between AXL and its ligands using surface plasmon resonance (SPR) as biochemical assay to investigate binding kinetics and functional digital spatial profiling (FuncOmap)^22^ to provide insights into the spatial interaction of AXL and its ligands in patient lung samples.

SPR analysis demonstrated that both rhGAS6 and rhPROS1 bind to rhAXL in a concentration dependent manner (**Figure 2A**). However, rhPROS1 required significantly higher concentrations to achieve a comparable binding response with rhAXL compared to rhGAS6 (**Figure 2B**). Notably, rhGAS6 exhibited a much higher binding affinity, with a dissociation constant (Kd) of 0.11 nM, compared to rhPROS1, which had a Kd of 52.15 nM. The purity of rhGAS6 and rhPROS1 was confirmed via SDS-PAGE (**Figure S2**).

**Figure 2.**
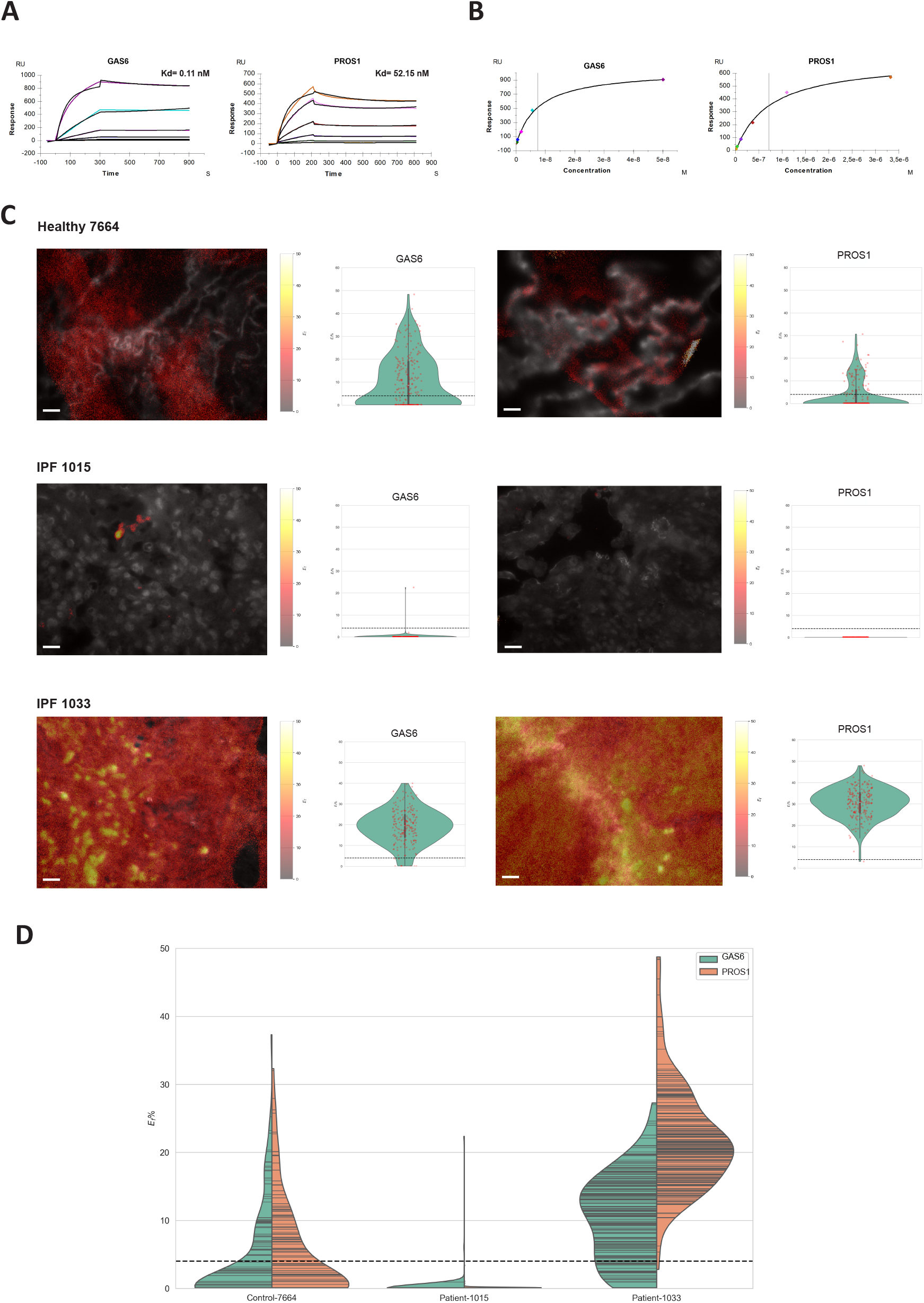
AXL interacts with both GAS6 and PROS1 ligands: determined by surface plasmon resonance and FuncOmap. **(A)** Surface plasmon resonance response curve showing the binding of rhGAS6 (Kd 0.11 nM) and rhPROS1 (Kd 52.15 nM) in different concentration to rhAXL. **(B)** Concentration dependent binding of rhGAS6 and rhPROS1 to rhAXL. Different colored lines in (A) correspond to different colored dots in (B), which indicate different concentrations of rhGAS6 and rhPROS1. **(C)** 7664 is a tissue section from a healthy lung whereas 1015 and 1033 are sections from IPF patients with 60% and 90% fibrosis, respectively. The panels show representative images mapping the interactive states of GAS6 and PROS1 with AXL (pseudo-color heat map of (Ef%) on the expression level of GAS6 and PROS1 (fluorescence images in grey scale). The individual violin plots quantify the heterogeneity of interactive states for both ligands and the receptor. The dotted line at 4% Ef represents the Förster Radius (R_0_). Values below 4% are not representative of ligand-receptor interactions. **(D)** Global violin plots of all coincident regions (per pixel) of AXL-GAS6 and AXL-PROS1. Each global violin plot represents 30x10^6^ data points. To determine the p values for AXL-GAS6 and AXL-PROS1, using non-parametric Mann-Whitney U test, we utilized 1000 randomly generated data points of the 30x10^6^. Healthy lung has similar distribution of AXL-GAS6 and AXL-PROS1 interactions, whereas the two patient samples show, in one case (1015) no interaction of AXL-GAS6 or AXL-PROS1 and another case (1033) a significantly higher distribution of AXL-PROS1 versus AXL-GAS6. The p values between AXL-GAS6 and AXL-PROS1 are 1.9x10^-3^ (7664), 2.4x10^-1^ (1015) and 8.2x10^-33^ respectively. The dotted line at 4% Ef represents the Förster Radius (R_0_). Scale bar represents 100 µm.

To further examine ligand-receptor interactions *in-situ*, we employed time-resolved Förster resonance energy transfer (TR-FRET) microscopy combined with FuncOmap^22^ . FuncOmap enables the automatic mapping of %Ef on each coincident pixel, identifying regions of interaction between AXL-GAS6 and AXL-PROS1. For these experiments, we analyzed tissue samples from healthy human lungs and IPF patients.

**Figure 2C** shows tissue sections from a healthy lung (7664) and from IPF patients (1015 and 1033) exhibiting varying degrees of disease progression. Patient 1015 exhibited 60% fibrosis, while patient 1033 showed more advanced fibrosis at 90% (**Figure S3**). The images display representative mappings of the interactive states of GAS6 and PROS1 with AXL, shown as pseudo-color heat maps of %Ef overlaid on grayscale fluorescence images representing GAS6 and PROS1 expression levels. Individual violin plots quantify the heterogeneity of ligand-receptor interactions for both GAS6 and PROS1 with AXL. The dotted line at 4% Ef represents the Förster Radius (R_0_), below which values are not indicative of ligand-receptor interactions.

**Figure 2D** provides a global violin plot summarizing all coincident regions of AXL-GAS6 and AXL-PROS1 interactions across tissue sections. Corresponding individual plots are shown in **Figure S4**. Each global violin plot represents 30x10^6^ data points. However, to determine a statistical significance using the p-values for AXL-GAS6 and AXL-PROS1, we applied the non-parametric Mann-Whitney U test to 1000 randomly generated data points.

Our analysis revealed that healthy lung tissue exhibits a similar distribution of AXL-GAS6 and AXL-PROS1 interactions. In contrast, the IPF patient samples show distinct patterns; sample 1015 demonstrates no detectable interaction between AXL-GAS6 or AXL-PROS1, while sample 1033 exhibits a significantly higher distribution of AXL-PROS1 interactions compared to AXL-GAS6. The calculated p-values between AXL-GAS6 and AXL-PROS1 are 1.9x10^-3^ (7664), 2.4x10^-1^ (1015) and 8.2x10^-33^ respectively. As IPF is a highly heterogeneous disease with patchy occurrence in the lung, we postulated that the difference in AXL interaction status in patients could reflect the different pathologies of IPF. To test this, we used a mouse model to further investigate the role of AXL in lung repair.

### Longitudinal analysis of lung repair processes after selective epithelial injury informs distinct phases of repair

Given the hypothesis that idiopathic pulmonary fibrosis (IPF) arises from repeated alveolar epithelial injury^2–4^, as well as the observed impact of AXL in the proliferation of basal cells, we sought to investigate mechanisms relevant for repair after epithelial injury, as changes in these mechanisms are thought to drive the early initiation phase of IPF. To achieve this, mice were intratracheally (i.t.) transduced with either AAV9-DTR or AAV6.2-DTR^23^ at a concentration of 10^11^ VG, to target alveolar cells (AAV9) or whole lung epithelial cells (AAV6.2), respectively. As a control, AAV6.2/9-stuffer, with non-coding “stuffer” DNA was utilized. To minimize the DT-off target effect described in our previous study^23^, as low as 50 ng DT (i.t.) was administered 14 days post AAV application to induce epithelial cell depletion. Repair was assessed at various time points (day 1, 2,4, 8, and 14) post DT. Body weight and the plasma surfactant protein D (SP-D) levels were monitored and quantified dynamically during the injury and repair process (**Figure 3A**). The preferential targeting of alveolar cells by AAV9 and whole lung epithelial cells by AAV6.2 was validated (**Figure S5**).

**Figure 3.**
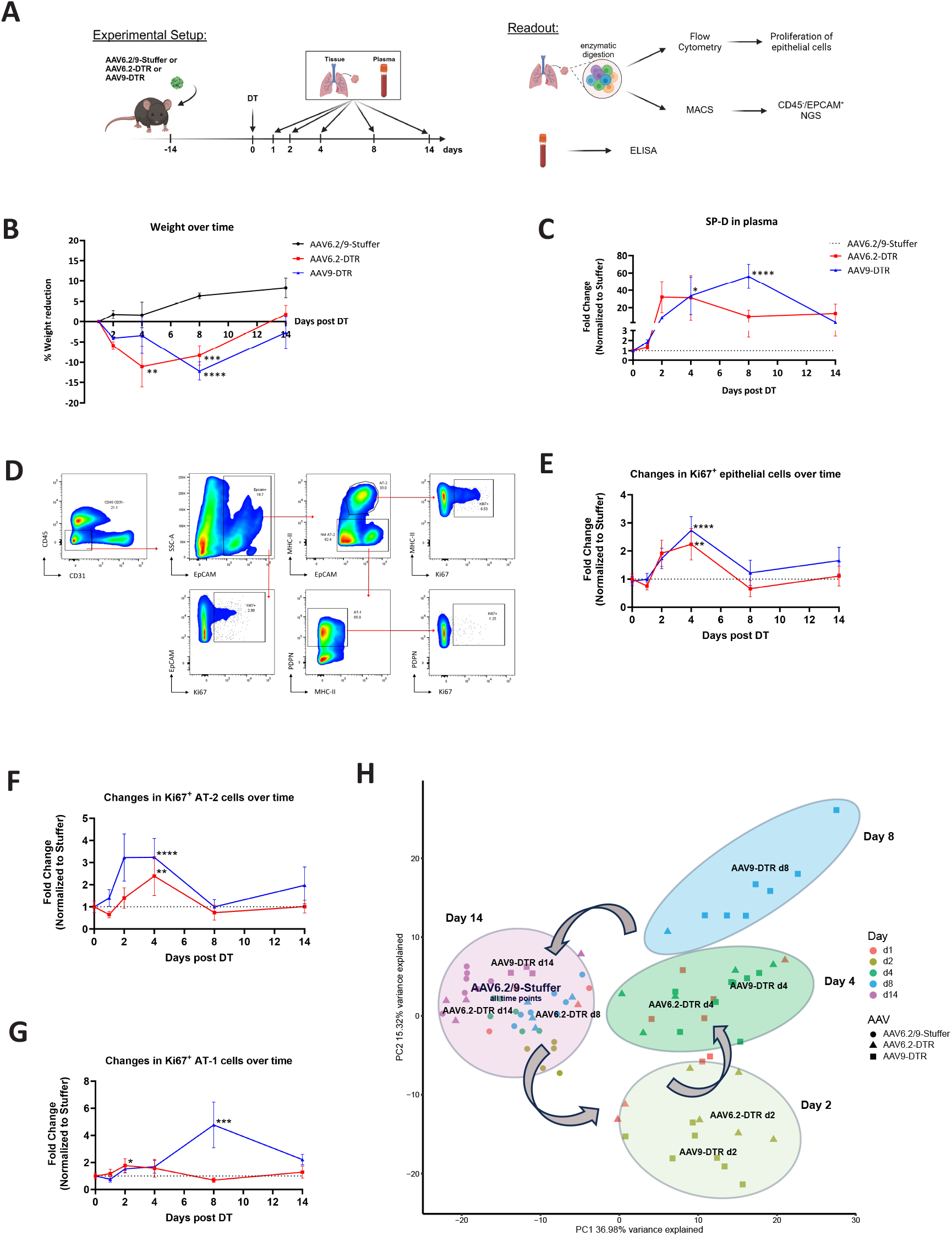
Longitudinal analysis of lung repair process after selective epithelial injury. **(A)** Experimental setup to study the normal lung repair mechanism after epithelial injury. AAV6.2/9-Stuffer/ AAV6.2-DTR/ AAV9-DTR were administered (i.t) into the mice. Lung tissue and plasma were collected at indicated time points post DT application for further readouts. Scheme was created in BioRender. Geillinger-kästle, K. (2025) https://BioRender.com/l78u806 **(B)** Weight of the mice over the course of repair. **(C)** Plasma SP-D level over the course of repair. Data were normalized to the corresponding stuffer control. **(D)** Representative flow cytometry gating to quantify epithelial cells proliferation. Cells were pregated for singlets. Proliferation of all epithelial cells: CD45^-^ CD31^-^ EpCAM^+^ Ki67^+^; AT-2 cells: CD45^-^ CD31^-^ EpCAM^+^ MHC-II^+^ Ki67^+^; AT-1 cells: CD45^-^ CD31^-^ EpCAM^+^ PDPN^+^ Ki67^+^. **(E-G)** Quantification of Ki67^+^ epithelial cells, AT-2 cells, and AT-1 cells. Data were normalized to the corresponding stuffer control. **(H)** PCA plot analysis of MACS-enriched epithelial cells (CD45^-^ EpCAM^+^) mRNA sequencing datasets. Data are shown as mean ± SEM of n= 3-7. P ≤ 0.05 (*); P ≤ 0.01 (**); P ≤ 0.001 (***); P ≤ 0.0001 (****) compared with stuffer control.

Weight loss was not observed in the stuffer group, instead, these mice gained weight. In contrast, significant weight loss occurred in the AAV6.2-DTR group at day 4 post-DT and in the AAV9-DTR group at day 8 post DT, with both groups recovering to baseline weight by day 14 (**Figure 3B**). Plasma SP-D levels, a biomarker for loss of epithelial barrier integrity, showed a corresponding increase, peaking at day 4 in the AAV6.2-DTR group (25-fold increase) and at day 8 in the AAV9-DTR group (50-fold increase) (**Figure 3C**). Plasma SP-D levels remained significantly elevated at day 8 for AAV9-DTR but not for AAV6.2-DTR, suggesting prolonged barrier leakage in the AAV9-DTR group. Based on this observation, we suggest that plasma SP-D levels mainly reflect alveolar leakage and to a lesser extent epithelial leakage.

Epithelial proliferation increased markedly in both models, with Ki67^+^ cells peaking at day 4 post-injury (**Figure 3D-E**), correlating with weight loss and elevated SP-D level (**Figure 3B-C**). Proliferation of alveolar type 2 (AT-2) cells began on day 2, peaking at day 4 post injury, with a higher induction observed in the AAV9-DTR group compared to AAV6.2-DTR (**Figure 3F**). In the AAV9-DTR group, Ki67^+^ alveolar type 1 (AT-1) cells increased at day 8 post injury, which may be attributed to the fact that Ki67 protein was not degraded directly after the proliferation. In contrast, only a minimal number of Ki67^+^ AT-1 cells were observed in the AAV6.2-DTR group (**Figure 3G**). By day 14 post DT, the Ki67^+^ cell count reverted to baseline in all groups (**Figure 3E-G**).

To further explore the molecular processes underlying lung epithelial repair, mRNA sequencing (mRNA-seq) was performed on magnetic activated cell sorting (MACS) enriched epithelial cells from all groups. Principal component analysis (PCA) of mRNA-seq data revealed that AAV6.2-DTR group from day 8 clustered together with both AAV6.2- and AAV9-DTR groups from day 14 and all stuffers groups from all-time points, whereas the remaining AAV-DTR samples formed a distinct cluster along PC1 (**Figure 3H**).

These results suggest that the AAV6.2-DTR injured lung has a faster repair rate in comparison to AAV9-DTR, with similar transcriptomic profiles suggesting repair in both groups by day 14 post injury as indicated by weight of the mice, SP-D level, Ki67^+^ cell count, and gene expression profiles.

### Upregulation of IPF signaling pathways involving *Axl* upon alveolar injury

To elucidate molecular mechanisms underlying lung injury and repair, Ingenuity Pathway Analysis (IPA) was performed on epithelial mRNA-seq data across the injury time course. Multiple canonical pathways were regulated, including the idiopathic pulmonary signaling pathway, which was predominantly driven by genes associated with epithelial-mesenchymal transition (EMT) (**Figure 4A**).

**Figure 4.**
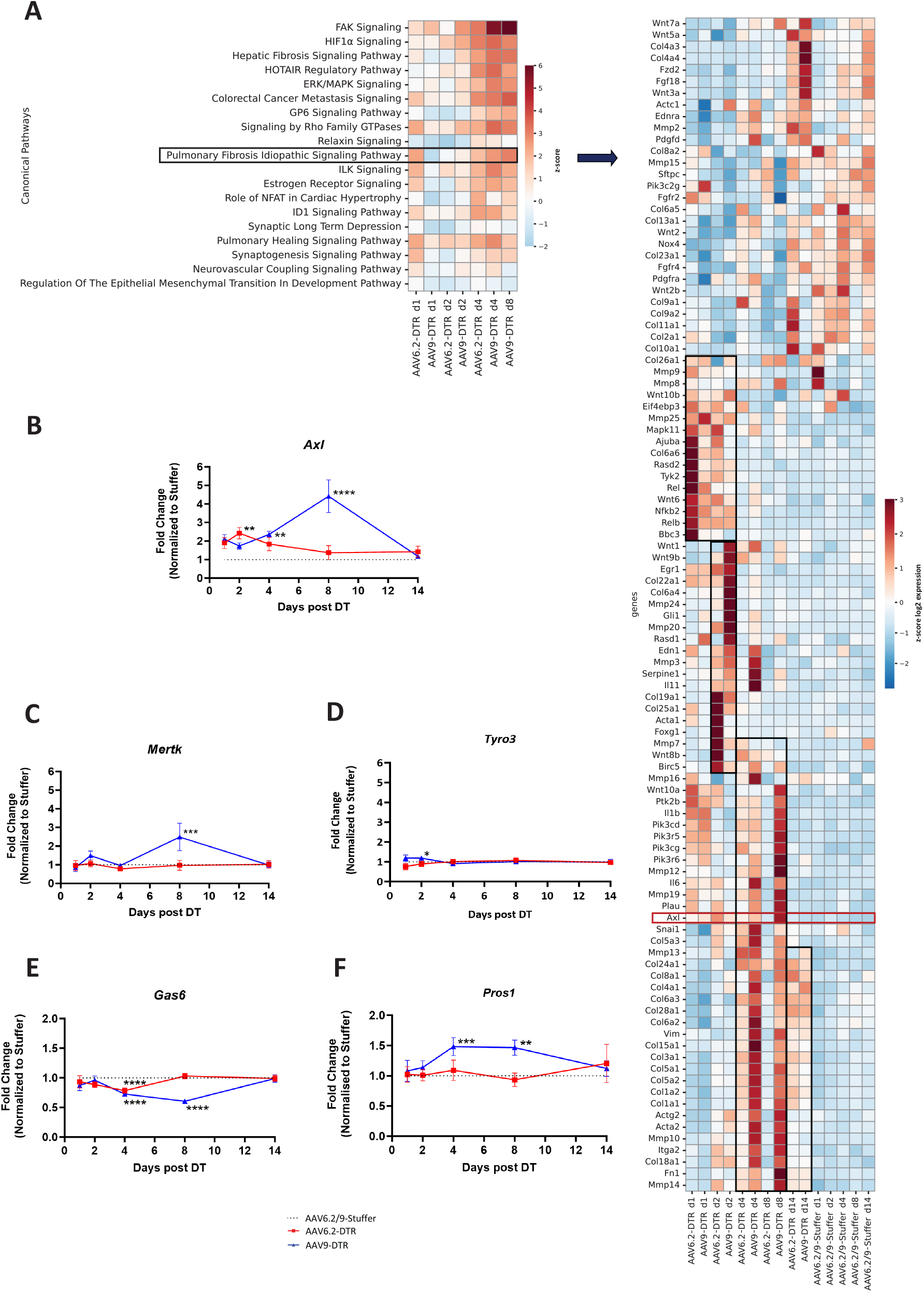
Upregulation of IPF signaling pathways involving *Axl* in response to alveolar injury. **(A)** IPF signaling pathway was one of the increasingly upregulated pathways over time as revealed by IPA analysis. The gene set contributing to this pathway included distinct groups of early and late profibrotic associated genes (marked by black rectangles) that were serially and time dependently upregulated. Timepoint order forced, genes hierarchical clustered. **(B-D)** Fold change of mRNA expression levels of TAM RTK family, *Axl, Mertk* and *Tyro3* over the course of repair. **(E-F)** Fold change of expression level of TAM RTK ligands, *Gas6* and *Pros1* over the course of repair. Data are shown as mean ± SEM of n= 3-7. P ≤ 0.05 (*); P ≤ 0.01 (**); P ≤ 0.001 (***); P ≤ 0.0001 (****) compared with stuffer control.

Hierarchical clustering of the genes along the time course revealed clusters of early and late profibrotic genes, indicating a sequential upregulation that coordinated the initiation of the repair process. Early response to lung injury (days 1 and 2 post DT) was characterized by acute inflammation and the upregulation of pro-inflammatory genes. By day 4 post DT, there was a transition from inflammation to early fibrotic signaling, evidenced by the increased expression of common fibrotic/ EMT markers (e.g. collagen family, *Acta2, Fn1, Vim*), which are the TGF-ß downstream genes^24^ . Additionally, *Axl* and its downstream signaling pathway, *Pi3k* were upregulated^25^. This coincided with the peak of epithelial cell proliferation (**Figure 3E**). On day 8 post DT, the expression of *Axl* and *Pi3k* isoforms were further enhanced, particularly in the AAV9-DTR group. By days 8 and 14, the downregulation of EMT associated markers compared to day 4 post DT suggests that repair was being achieved (**Figure 4A**).

Days 4 and 8 emerged as critical phases of repair, corresponding to maximal proliferation and subsequent differentiation, respectively **(Figure 3E)**. During this crucial period, *Axl* was specifically upregulated following alveolar-specific injury (**Figure 4A-B**), whereas *Mertk* only showed upregulation on day 8 in AAV9-DTR group (**Figure 4C**), and *Tyro3* remained unchanged (**Figure 4D**). Analysis of AXL ligands revealed downregulation of *Gas6* and upregulation of *Pros1* during days 4-8, with both returning to baseline by day 14 (**Figure 4E-F**).

Together, these data suggest that epithelial repair requires a ligand-specific shift in AXL signaling, characterized by Gas6 downregulation and Pros1 upregulation to terminate the proliferative phase. Disruption of this balance, as observed in IPF where both ligands remain elevated, may impair normal repair and promote fibrotic progression.

### Quantitative functional imaging to investigate AXL-GAS6/PROS1 interactive states under stress repair

Under normal repair conditions in the AAV-DTR model, alveolar-specific injury induced upregulation of *Axl* and *Pros1* and downregulation of *Gas6* compared to stuffer controls at days 4 and 8 post injury (**Figure 4E-F**).

Given that smokers constitute a large proportion of IPF patients^2,4,16^, we mimicked stress repair in IPF by exposing mice to chronic cigarette smoke for 28 days prior to AAV-DTR injury (**Figure 5A**). Day 4 post injury, AXL-ligand interactions were assessed *in-situ* using TR-FRET. Representative images reveal heterogenous AXL-ligand interaction states within individual mouse (**Figure 5B-C)**. In one representative lung, distinct coincident regions exhibited low AXL-GAS6 interactions (%Ef of 1.55% and 6.96%), compared with markedly higher AXL-PROS1 interactions (%Ef of 14.34% and 19.09%).

**Figure 5.**
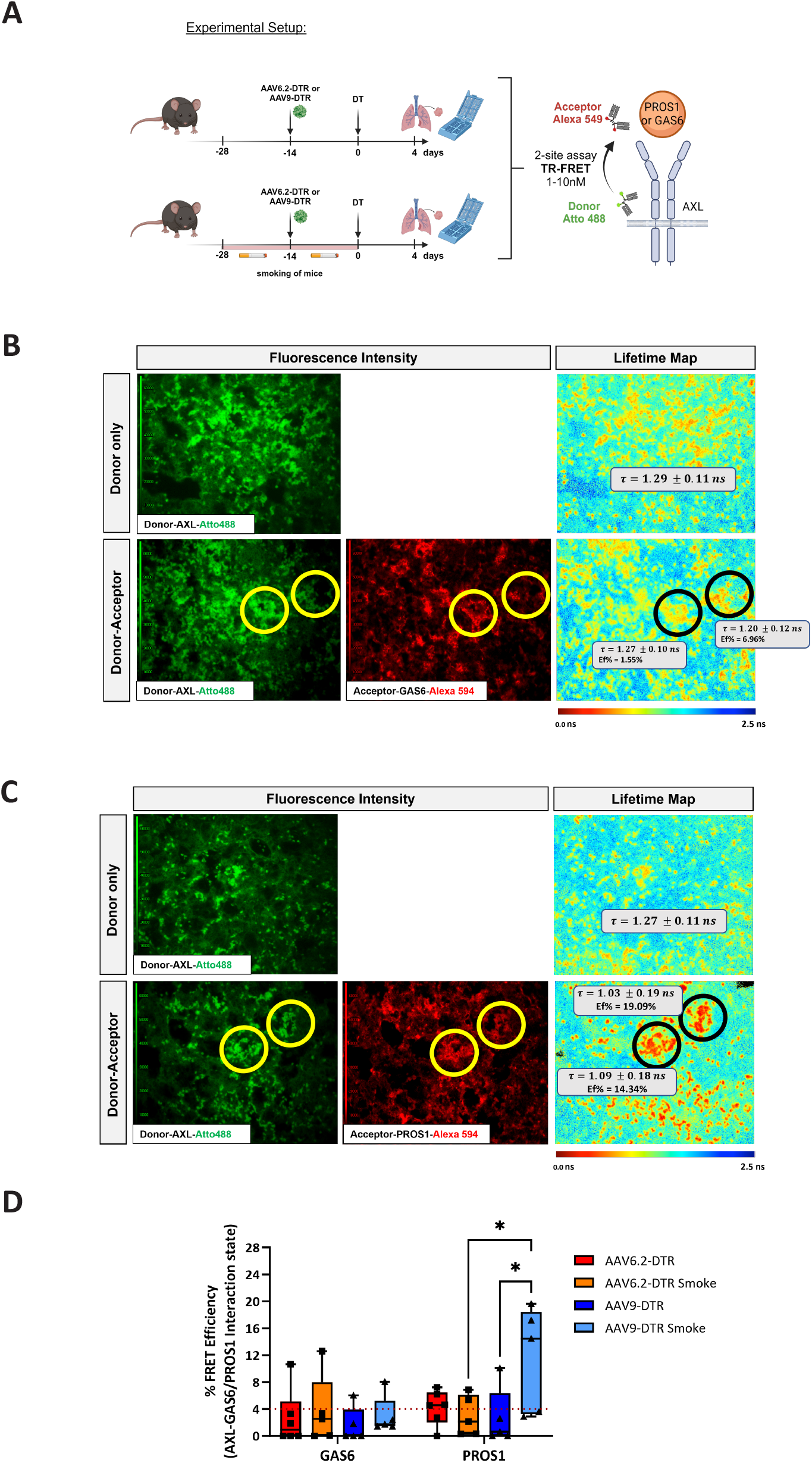
Enhanced AXL-PROS1 interactions following alveolar injury under stress repair conditions. **(A)** Schematic of the experimental setup to study AXL-GAS6/PROS1 interaction using aFRET in mouse lung tissue sections. Lungs were isolated on day 4 post DT. aFRET is a coincidence assay, labels both receptor (AXL) and ligands (GAS6 or PROS1) (see Methods for details). FRET occurs in the range of 1-10 nm. Scheme was created in BioRender. Geillinger-kästle, K. (2025) https://BioRender.com/h45x771 **(B)** Representative image of AXL-GAS6 interaction. **(C)** Representative image of AXL-PROS1 interaction. **(B-C)** Upper figures showed the mean of the donor lifetime (𝒯) in the absence of acceptor. Lower figures showed the interaction state between AXL-GAS6/ PROS1 for each coincidental region (marked by yellow and black circles). Different coincidental regions possess different interaction state. **(D)** Box and whiskers plots showed the heterogeneity of AXL-GAS6/PROS1 interaction states represented as FRET efficiency (Ef%). Ef% below 4% represents the Förster Radius (R_0_ at 5.83nm) and not proteins, interactive state as marked by red dashed line. Data are shown as box and whiskers plot of n=5-6. Each dot represents median of Ef% for each mouse. P ≤ 0.05 (*).

For each individual mouse, 10 coincident regions were analyzed, and the median of the %Ef of all acquired coincident areas are represented as box and whisker plots. Notably, a higher interactive state between AXL and PROS1 was observed in the AAV9-DTR smoke-treated group compared to its non-smoke counterpart and the AAV6.2-DTR smoke group (**Figure 5D**).

These findings suggest that AXL-PROS1 interactions are enhanced following alveolar injury under stress repair conditions, such as those proposed in IPF.

## DISCUSSION

IPF is a progressive lung disease characterized by aberrant epithelial repair and fibrosis, yet its early pathogenesis remains poorly understood due to reliance on late-stage biopsies. To address limited insight into early IPF pathogenesis, we used human data and mouse models to elucidate the role of AXL and its ligands GAS6 and PROS1 in epithelial repair and fibrosis.

Single-cell RNA sequencing revealed AXL upregulation in basal and aberrant basaloid cells in IPF lungs, cells which are uniquely enriched in IPF lungs^5–7^. In contrast, GAS6 and PROS1 were enriched in aberrant basaloid- and AT-1 cells (**Figure 1A**). Consistently, BALF analysis from IPF patients demonstrated elevated GAS6 and PROS1 levels, with PROS1 being significantly higher than GAS6, highlighting the role of AXL signaling in epithelial dysfunction in IPF (**Figure 1B**).

Given AXL’s predominant expression in basal cells, we employed primary human small airway epithelial basal cells (SAECs) to explore its functional role in epithelial repair. Our data demonstrate that AXL expression correlates strongly with epithelial proliferation (**Figure 1G**), an effect primarily driven by GAS6, consistent with previous studies^43^. Interestingly, while PROS1 alone did not significantly alter epithelial behavior, it attenuated GAS6-mediated proliferation, suggesting a shift from a proliferative to a differentiation phase. Supporting this, AXL knockdown in human lung multipotent cells promoted an AT-2 phenotype^38^, reinforcing the hypothesis that PROS1 modulates GAS6-driven AXL signaling to facilitate proper epithelial repair.

Historically, GAS6 has been considered the primary ligand of AXL^8,9^. However, we have confirmed the direct interactions between AXL and both GAS6 and PROS1 using surface plasmon resonance (SPR) and time-resolved Förster resonance energy transfer (TR-FRET) microscopy. Spatial mapping of ligand-receptor interactions in human lung tissue revealed distinct patterns in IPF patients highlighting the heterogeneity of this disease, with a higher prevalence of AXL-PROS1 interactions compared to AXL-GAS6 in a donor with more advanced disease stages (**Figure 2**), as well as in a mouse model of alveolar injury under stress repair conditions (**Figure 5**). These findings underscore the importance of ligand-specific regulation in AXL signaling during repair processes.

Interestingly, despite the significantly higher Kd for PROS1 binding to AXL compared to GAS6 (**Figure 2A**), high concentration of PROS1 was able to fully blunt the GAS6 effect on proliferation (**Figure 1H**). This discrepancy may reflect limitations of the reductionist *in-vitro* approach used in SPR to determine Kd values, which does not fully capture the complexity of the cellular/ molecular environment. Factors such as molecular crowding, membrane compartment dynamics, and spatial organization impact ligand-receptor interactions and downstream signaling. Thus, *in-situ* Kd values do not necessarily reflect in vitro Kds. Furthermore, alternative signaling pathways may contribute to AXL signaling modulation by PROS1. While these pathways remain to be fully elucidated, they do not exclude the involvement of PROS1 in modulating AXL signaling.

Since IPF is proposed to be caused by aberrant alveolar repair, this study utilized an AAV9-DTR/DT mouse model to specifically injure alveolar cells and study the subsequent repair process, as opposed to whole lung epithelium injury mediated by AAV6.2-DTR/DT. This model allowed us to mimic the early initiation phase of alveolar epithelial injury observed in IPF and provided a more time- and cost-effective method for targeting specific cell types compared to the generation of transgenic mouse models.

Temporal analysis of the repair process revealed distinct phases, with epithelial proliferation peaking at day 4 post-injury and differentiation occurring at day 8 (**Figure 3E**). IPA analysis of epithelial cell datasets revealed the temporal regulation of gene sets involved in IPF signaling pathways, suggesting a coordinated gene expression pattern essential for initiating homeostatic repair (**Figure 4A**). Disruption of these patterns may lead to aberrant repair and fibrosis. Interestingly, during this critical repair window, AXL signaling was among the pathways that exhibited this specific pattern, particularly following alveolar-specific injury. While AXL itself was highly upregulated, its ligands GAS6 and PROS1, displayed opposing expression patterns, with GAS6 being downregulated and PROS1 being increased, suggesting a ligand-specific shift in AXL signaling during normal repair, as this mouse model repairs completely at day 14 post injury indicated by weight of the mice, SP-D level, Ki67^+^ cell count, and gene expression profiles (**Figure 3A-H**). This switch from GAS6 to PROS1 may be essential for shifting from proliferation to differentiation, a balance that appears disrupted in IPF.

The differential expression of AXL and its ligands as well as its molecular interactions with both its ligands, along with the upregulation of EMT-associated genes, suggests that targeting AXL signaling could modulate fibrotic responses and improve repair outcomes. The complexity of ligand interaction with AXL underscores the need for a balance between GAS6 and PROS1 to facilitate normal lung repair, highlighting the necessity for precise therapeutic targeting in IPF. Since AXL is predominantly expressed in basal cells, excessive AXL-GAS6 interaction may lead to an uncontrolled basal cell proliferation, which in turn could result in insufficient differentiation into AT-2 cells.

The delicate balance of proliferation, differentiation, and apoptosis of AT-2 cells is essential in successful lung repair^2,26,27^. In IPF, this balance is probably severely disturbed, due to inefficient and insufficient differentiation of basal cells into AT-2 cells, leading to the loss of AT-1 cells, impaired repair, and progression into fibrosis. This imbalance may explain the abnormal expansion of basal cells into the distal parts of the lung (bronchiolization) in IPF^7,28^ . Our refined model (**Figure 6**) proposes a “Yin-Yang” relationship between PROS1 and GAS6, which appears to be disrupted in IPF patients. In these patients, although PROS1 is upregulated as a repair effort, the concurrent upregulation of GAS6 hinders the normal repair process.

**Figure 6.**
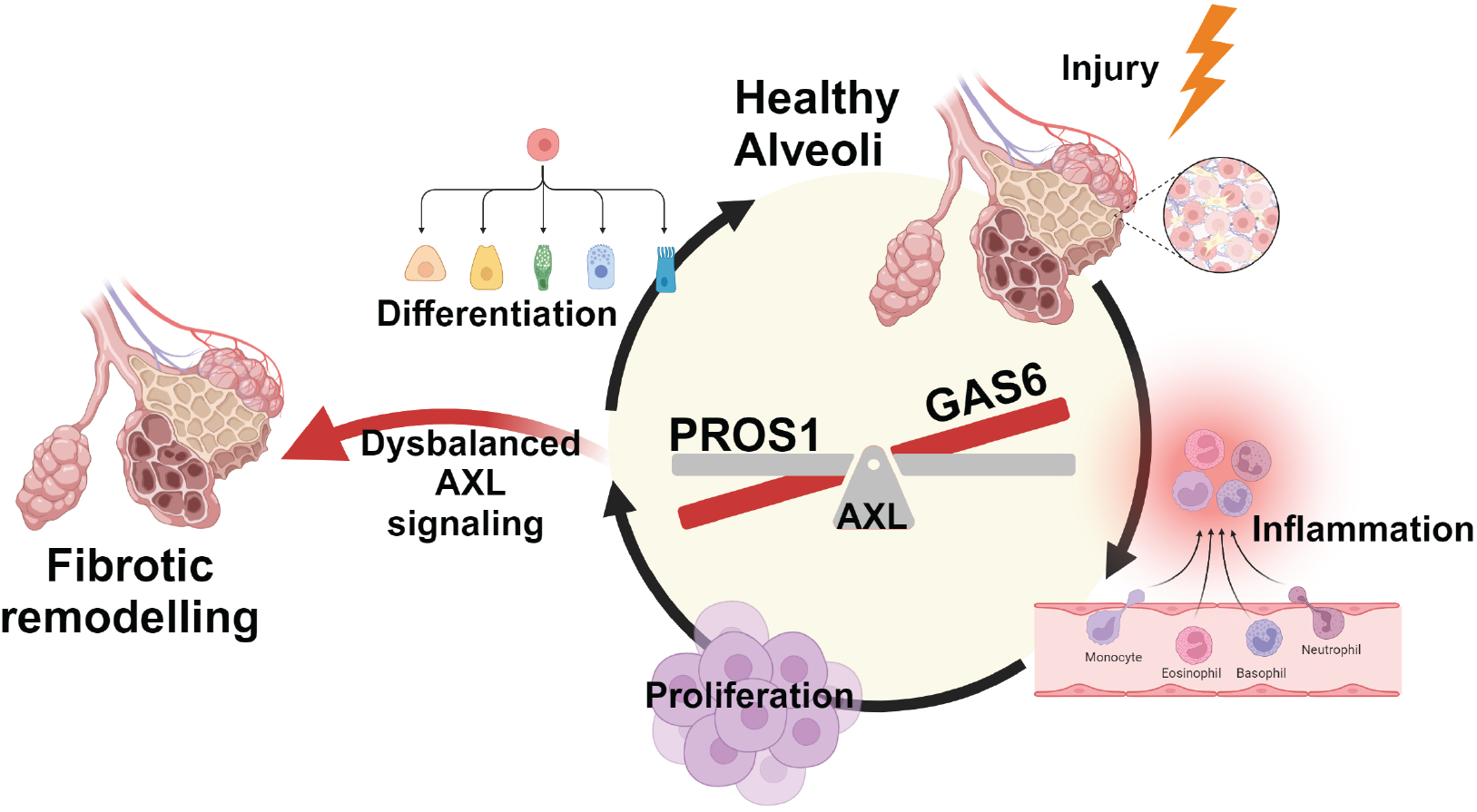
Refined mechanism of AXL-GAS6/PROS1 interaction during lung repair. After injury, the normal repair process involves an initial inflammatory response, followed by AXL-GAS6 interaction, which promotes cell proliferation. Once sufficient proliferation is achieved, AXL-PROS1 binding occurs, halting proliferation and triggering cell differentiation. This balance between proliferation and differentiation ensures proper repair, leading to healthy alveoli. However, dysbalanced AXL signaling due to repeated injury and persistently high TGF- ß levels, can result in aberrant repair and irreversible fibrosis (IPF). This may occur due to high-affinity interaction between AXL and GAS6 prevents PROS1 from effectively binding to AXL. As a result, excessive proliferation occurs with insufficient differentiation, leading to aberrant repair and fibrosis. Scheme was created in BioRender. Geillinger-kästle, K. (2025) https://BioRender.com/q77c168.

In conclusion, we have demonstrated for the first time via FuncOmap, the direct in situ and spatial interactive states between AXL and PROS1. BALF analysis from IPF patients and TGF-ß stimulated SAEC revealed significantly higher PROS1 levels compared to GAS6, supported by the spatial molecular showing higher AXL-PROS1 interactions compared to AXL-GAS6 in a donor with more advanced disease stage, underscoring its biological relevance. These findings have important implications not only for IPF treatment, but also in the cancer field, where GAS6-AXL signaling in immune cells is already being targeted. Our data show that AXL signaling in disease is not solely mediated by GAS6, rather the balance between PROS1/GAS6 and AXL plays a pivotal moment in directing healthy versus aberrant repair.

## METHODS

### Human bronchoalveolar lavage fluid (BALF)

Human non-diseased and COPD BALF samples were purchased from Tissue Solutions, Ltd., UK and ILD/IPF BALF samples were received from Wangen hospital, Germany. Patients consent was obtained for the use of samples in research applications. Information on BALF donors is provided in **Supplementary Table S1**.

### SAEC submerged culture

T175 flasks were coated with 30µg/ml rat tail collagen (Corning, #354236) for 45 minutes. SAEC from 7 healthy human donors (Lonza) were thawed into precoated flasks containing SAEC submerged complete medium (490 ml Pneumacult™ Ex Plus basal medium (STEMCELL, #05041) supplemented with 10 ml 50x Pneumacult™ Ex Plus supplement (STEMCELL, #05042), 0.5 ml hydrocortisone (STEMCELL, #07925), and 5 ml penicillin-streptomycin (ThermoFisher, #15140122). At 80% confluence, cells were detached using ACF enzymatic dissociation kit (STEMCELL, #05426), centrifuged at 300g for 5 minutes. 50,000 cells were then seeded into 24-well plates. The cells were then serum-starved in additive-free DMEM (ThermoFisher, #11966025) for further assays.

### SAEC air liquid interface (ALI) culture

SAEC were thawed and detached from the flask using the same method as described previously. After centrifugation, 30,000 cells were seeded onto the rat tail collagen coated transwell insert using complete SAEC submerged medium. The medium was added to the apical and basolateral chambers of the wells until they reached confluence. Once confluence was achieved, the cells underwent airlifting by removing the apical media entirely and replacing the basal media with SAEC ALI complete medium (450 ml Pneumacult™ ALI-S medium (STEMCELL, #05051) supplemented with 50 ml 10x ALI-S supplement (STEMCELL, #05052), 2.5 ml hydrocortisone, 5 ml maintenance system supplement (STEMCELL, #050502), 1 ml heparin (STEMCELL, #07980), and 5 ml penicilin-streptomycin) to promote cell differentiation. Subsequent assays were conducted in SAEC ALI complete medium. Cells are fully differentiated 21 – 30 days post ALI.

### Generation of CRISPR-Cas9 AXL knockout SAEC

A ribonuclear protein (RNP) complex was assembled by incubating 1250 ng of TrueCut™ Cas9 protein V2 (ThermoFisher, #A36498) with 7.5 pmol TrueGuide™ Mod human AXL sgRNA (ThermoFisher, CRISPR932418_SGM) for 15 – 30 minutes at RT. Meanwhile, 6- or 24-well plates were coated with rat tail collagen and filled with antibiotic-free SAEC submerged complete medium. Electroporation was performed using the AMAXA Basic Nucleofector Kit for Primary Mammalian Epithelial Cells (Lonza, #VPI-1005) following the manufacturer’s protocol. Briefly, 2 µg of pmaxGFP™ vector was added to the nucleofection cuvette, followed by 700,000 cells resuspended in 100 µl nucleofector solution and mixed with the RNP complex. The mixture was electroporated using program T-020, after which 500 µl pre-warmed medium was added and cells were immediately transferred to prepared plates. Cells were ready for downstream assays 24 – 48 hours post-electroporation.

### Bromodeoxyuridine (BrdU) proliferation assay

50,000 SAEC were cultured in a 24-well plate with SAEC submerged complete medium. The following day, cells were starved overnight in DMEM. Cells were then treated with rhGAS6 (400 ng/ml), rhPROS1 (1200 ng/ml), or combination of both for 48 hours. Bromodeoxyuridine (BrdU) assay was performed according to the manufacturer’s protocol (Roche, #11647229001). Briefly, cells were labelled with BrdU for 4 hours, fixed, and incubated with anti-BrdU antibody, followed by substrate addition. Optical density was measured using a SpectraMax microplate reader (Molecular Devices) at 370 nm with a reference wavelength of 492 nm.

### Two-site amplified Time Resolved-Forster Resonance Energy Transfer (aFRET)

Paraffin embedded human and mouse lungs (4 µm) sections were subjected to dewaxing and rehydration, followed by heat antigen retrieval. After peroxidase suppression and blocking with 3% BSA in PBS, sections were incubated overnight at 4°C with primary antibodies (Donor: AXL (1:100) (Abcam, #ab130218 or #AF854), Acceptor: GAS6 (1:100) (Cell signaling, #67202S or ThermoFisher, #BS-7549R)) or PROS1 (1:100) (Sigma-Aldrich, #HPA007724 or ThermoFisher, #14-5791-85) in 1% BSA in PBS. For secondary antibody labelling, slides designated as donor-only were incubated with self-conjugated F(ab’)_2_-Atto488 (Jackson ImmunoResearch, #715-006-150 or Jackson ImmunoResearch #705-006-147, Sigma Aldrich, #41698-1MG-F), while those labelled as donor-acceptor were exposed to F(ab’)_2_-Atto488 and F(ab’)_2_-HRP (Jackson ImmunoResearch #711-036-152) for 2 hours at RT. Tyramide signal amplification was conducted on donor-acceptor slides, where F(ab’)_2_-HRP was bound to the acceptor primary antibody. Tyramide, conjugated to the Alexa594 chromophore (ThermoFisher, #B40925), was introduce to the HRP molecule, thereby fluorescently labelling the acceptor site. Finally, the slides were mounted using prolong diamond anti-fade mounting medium. This protocol was adapted from Veeriah, *et al*. (2014)^29^.

This two-site assay (aFRET) is patented: Patent US 10578,620 B2: Methods for detecting molecules in a sample: patent rights are held by The Francis Crick Institute.

### aFRET determined by frequency-domain Fluorescence Lifetime Imaging

Quantitative molecular imaging was performed using a custom-built, semi-automated frequency-domain fluorescence lifetime imaging (FLIM) system based on a modified Lambert FLIM. Donor and donor-acceptor samples are excited at 473 nm using a solid-state diode laser in a homodyne configuration modulated at 40 MHz.

The reduction in donor lifetime due to resonance energy transfer, caused by the acceptor, correlated with measurement of distances in the range of 1-10 nm allows the quantification of receptor-ligand interactions. Coincidence regions, where both donor and acceptor signals were identified and a total of 10 regions of interest (ROIs) selected within those regions. The fluorescence lifetimes and standard deviations are automatically calculated and exported for further analysis.

### Photophysical parameters for quantification of protein interactive states

Donor lifetime image in the presence (**τ**_**DA**_) and absence **(τ**_**D**_) of acceptor were automatically calculated, and FRET-efficiency (%Ef) was derived from the relative reduction in donor lifetime:

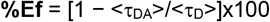

The Förster radius (R_0_) for Atto-488/Alexa 594 pair is 5.83nm, corresponding to 50% energy transfer. R_0_ of 5.83nm corresponds to 4% Ef, therefore an Ef lower than 4% is considered non-interactive.

### Functional Digital Spatial Profiling (FuncOmap) of AXL and Its Ligands GAS6 and PROS1

Functional mapping of coincident regions from the two-site assay was performed as described by Safrygina, *et al*. 2024^22^ . Briefly, the initial data capture from the FLIM images provides a donor and a donor-acceptor image. For each pixel the lifetime is recorded in the control donor and the donor-acceptor image and hence the FRET efficiency calculated automatically. The FLIM output is processed using the FuncOmap software^22^, outputting a spatially resolved FRET efficiency map, where each pixel value corresponds to the calculated FRET efficiency. A heatmap is applied to the FRET efficiency at a scale of 0-50% across the region of interest. The %Ef is mapped automatically on the acceptor fluorescent image (grey scale).

FuncOmap software is available under license from the University of Bath. For enquiries, contact Julian Padget and/or Banafshé Larijani.

### Analysis of the Functional Spatial mapping

The per pixel FRET efficiency values from the maps are plotted automatically as violin plots. For each mapped coincidence region there is a unique violin plot showing the distribution of the interactive states of AXL-GAS6 and AXL-PROS1. Each global violin plot represents 2x10^6^ data points. To determine p values for AXL-GAS6 and AXL-PROS1, we used the non-parametric Mann-Whitney U test with 1000 randomly selected data points from the 2x10^6^.

### Surface plasmon resonance to study AXL-GAS6/PROS1 binding kinetics

Binding kinetics of recombinant human GAS6 (rhGAS6) (R&D, #885-GSB) and recombinant human PROS1 (rhPROS1) (R&D, #9489-PS) to recombinant human AXL was assessed using surface plasmon resonance using a Biacore-T200 system. Biotinylated AXL (Boehringer-Ingelheim, #33-227) was immobilized on a streptavidin chip. Increasing concentrations of rhGAS6 (0.000069 µM - 0,05 µM; 1:3) or rhPROS1 (0.00069 µM - 5 µM; 1:3) were then injected onto the immobilized target in 25mM MES, 150 mM NaCl, 10 mM CaCl2, 0.05% Tween, pH 6.0. Mean association rate (K_on_) and dissociation rate (K_off_) was calculated from 3 individual experiments using Biacore-T200 analysis software. The dissociation constant (Kd) was calculated from K_on_/K_off_.

### Animals

Male C57BL/6JRj mice (8-12 weeks old) were purchased from Janvier Labs or Charles River and housed in individually ventilated cages at 22-25 °C, 12 h light cycle, and *ad libitum* access to food and water. Animal procedures were approved by Regierungspräsidium Tübingen, Germany (TVV-20-006-G, TVV 18-030-O).

### AAV to overexpress GFP or human DTR

Recombinant AAVs were produced following the procedure described in Strobel, *et al*. 2019^30^. Following sterile filtration, AAV preparations were titrated by digital PCR, using an ITR-specific primer/probe set. Self-complementary CMV-eGFP-poly(A) or CMV-hDTR-p(A)^23^ expression cassettes were packaged into either AAV6.2 or AAV9.

### Statistical analysis

The data are represented as means ± standard error mean (SEM). Statistical analysis was performed using GraphPad Prism. Mean values were compared using t-test for experiment with two groups and one-way- or two-way ANOVA for experiments with three or more groups, followed by Tukey’s or Dunnet’s multiple comparisons test. Median values of FRET efficiencies were compared in box and whiskers plots and the global violin plots from FuncOmap were compared using non-parametric Mann-Whitney U test. Correlation analysis was performed using Pearson or Spearman. p-values are represented as asterisk with, p > 0.05 (ns/ non-significant); p ≤ 0.05 (*); p ≤ 0.01 (**); p ≤ 0.001 (***); p ≤ 0.0001 (****).

## Acknowledgements

We thank Sylvia Blum, Anita Schönleber, Annika Meier, Michael Schilling, Eva Thaler, and Helene Lichius for assistance with the *in-vivo* experiments, Birgit Stierstorfer, Tanja Schönberger, Fabian Heinemann, and Nadine Rehm for their support in histology, Wioletta Skronska-Wasek and Wangen Hospital, Germany for the human IPF BALF samples. Marc Kästle for providing SAEC mRNA seq data. Holger Schlüter and Medizinische Hochschule Hannover (Hannover medical school) for providing the human lung sections. This work is funded by Boehringer Ingelheim Pharma GmbH & Co. KG, Germany. Schemes were created in BioRender.

## Author contributions

DS designed and performed most of the experiments and analyzed data. DS wrote the manuscript. CV and AD supervised sequencing experiments. DSc and WR performed sequencing experiments. CHM, KFC, and FR analyzed sequencing data. AF performed experiments. GS and YH supervised and performed SPR experiments. DS and CJA supervised and performed TR-FRET experiments. JP, SL, and BL performed FuncOmap analysis and interpretation of functional spatial mapping. BS provided AAV and conceptual advice. MJT provided conceptual advice. SGW, BL, and KGK conceptualized, supervised the study, and designed experiments. All authors edited and commented on the manuscript.

## Data availability

RNA-seq data generated within this study is deposited in the Gene Expression Omnibus (GEO) and will be made accessible upon acceptance of the manuscript. All other data is available upon request.

