## Supplementary Files for "AXL-GAS6/PROS1 Interaction: A Critical Switch Between Aberrant- and Healthy Repair Following Alveolar Lung Injury"

### Supplementary Figures

## S1

A

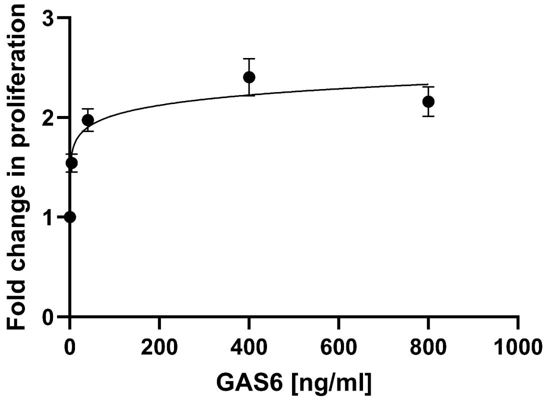


B

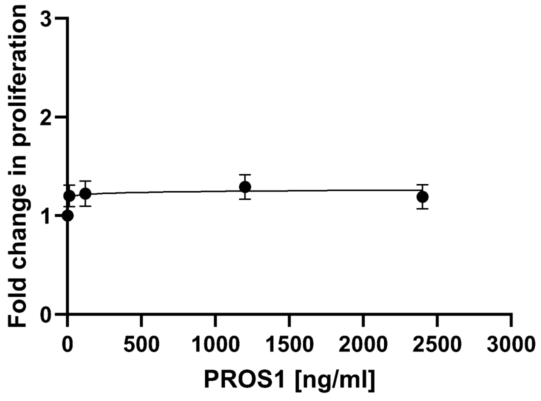


## S2

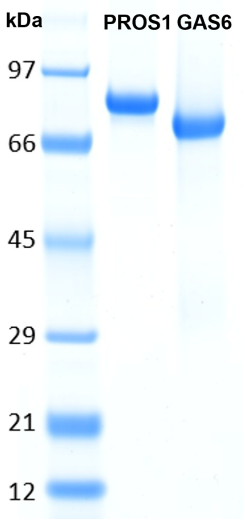


## S3

A Healthy 7664

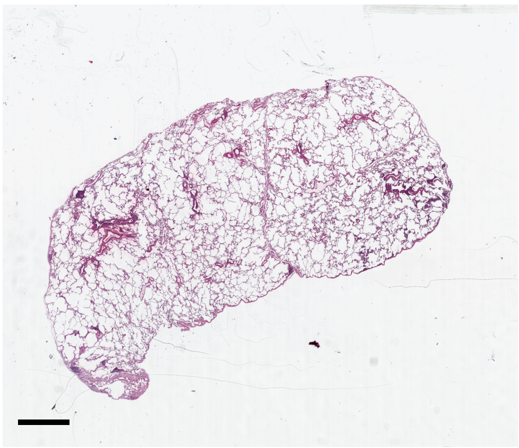

B IPF 1015

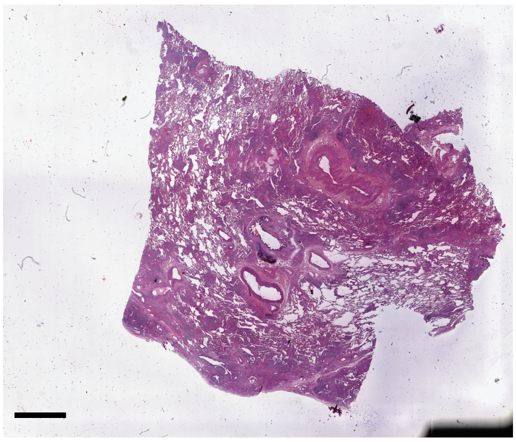

C IPF 1033

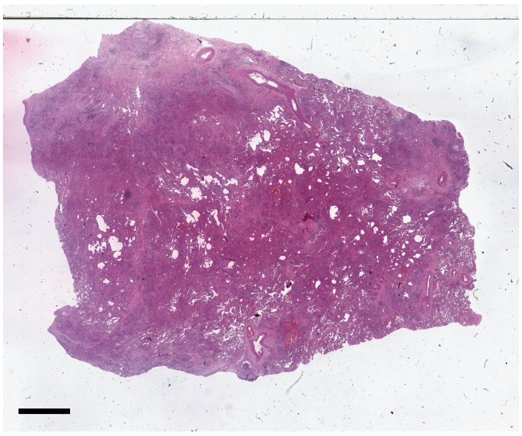

# S4

#### A Healthy 7664

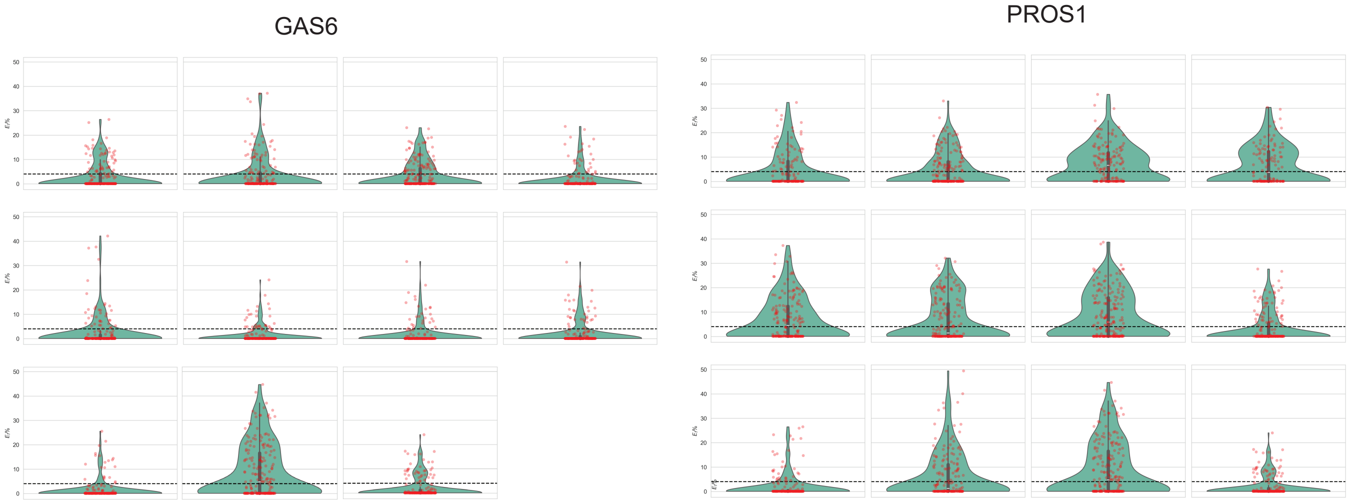


#### B IPF 1015

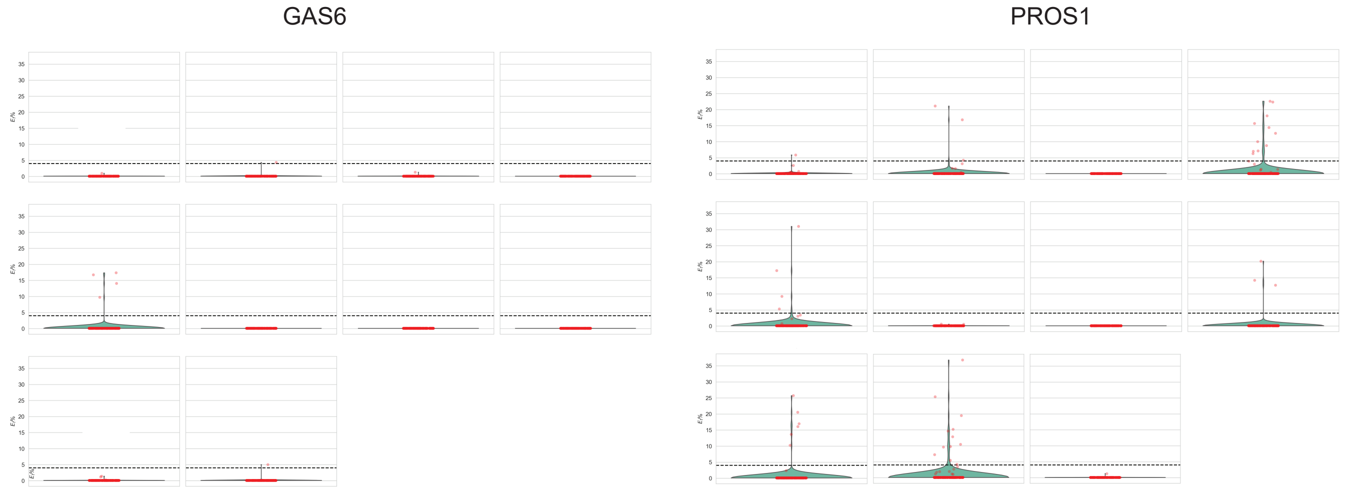


#### C IPF 1033

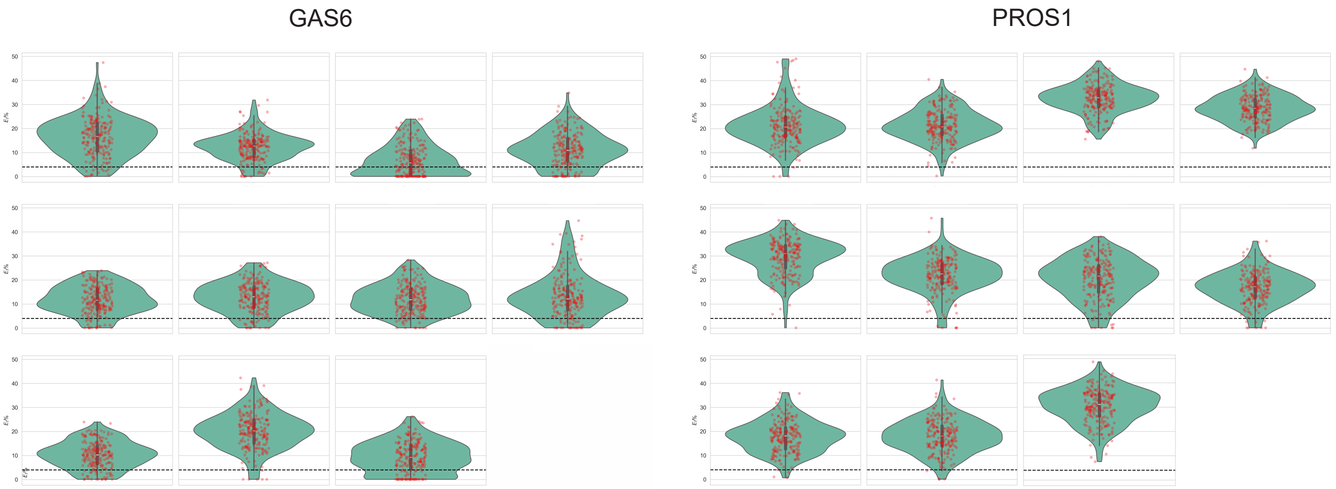

# S5

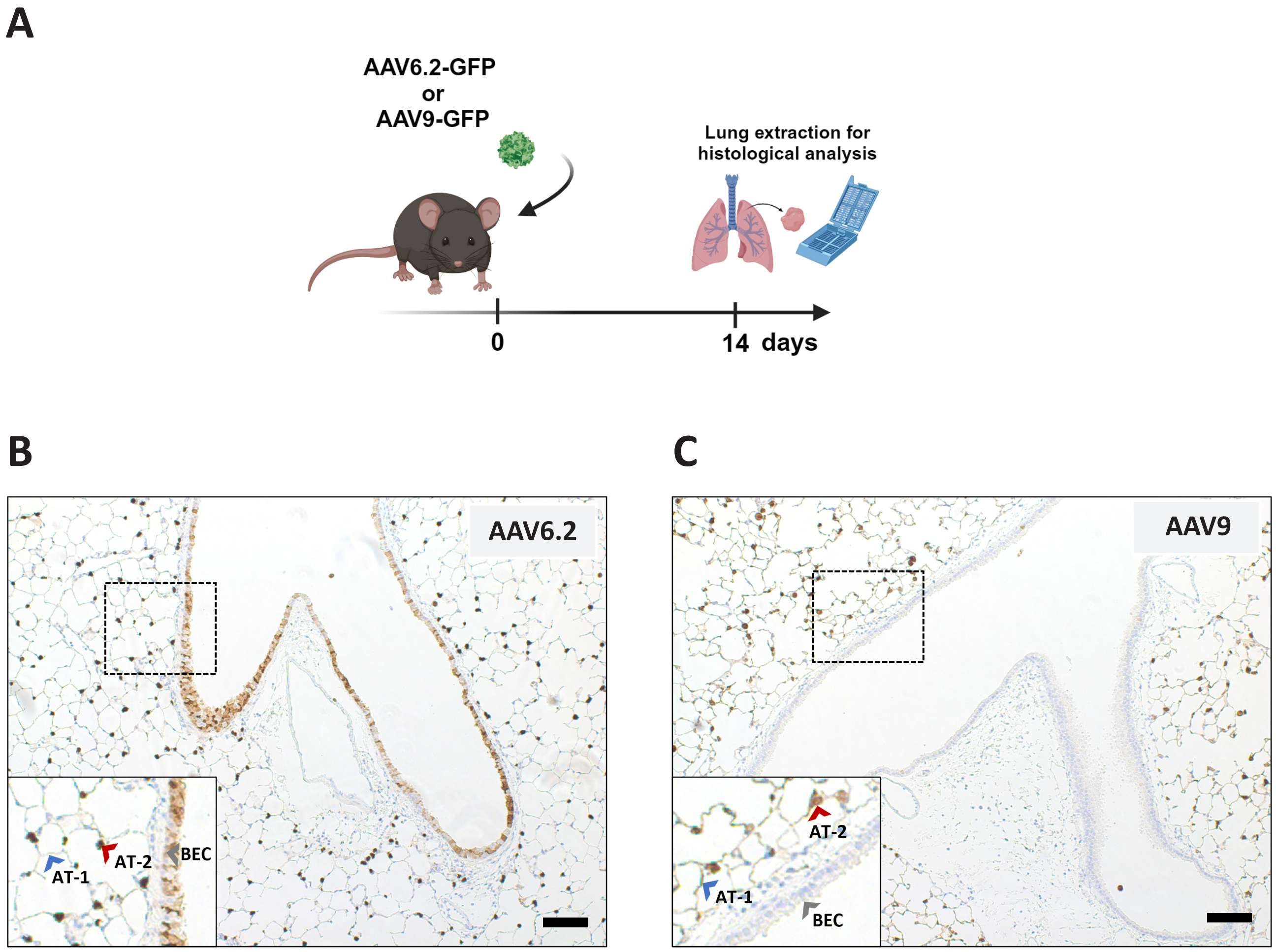
A
AAV6.2-GFP
or
AAV9-GFP
Lung extraction for
histological analysis
0
14 days
B
AAV6.2
AT-1
AT-2
BEC
C
AAV9
AT-2
AT-1
BEC

**Figure S1 Dose-dependent relationship between rhGAS6 or rhPROS1 and SAEC proliferation.** **(A)** rhGAS6 and **(B)** rhPROS1 dose influenced the proliferation of SAEC basal cells as measured by BrdU. 400 ng/ml rhGAS6 and 1200 ng/ml rhPROS1 were selected for all cell culture experiments. Data are shown as mean ± SEM of n= 6.

**Figure S2 Evaluation of rhPROS1 and rhGAS6 protein purity prior to SPR analysis.** SDS-PAGE proved the purity of rhPROS1 and rhGAS6 at expected molecular weight ~80 kDa and ~70 kDa, respectively.

**Figure S3 H&E images of human lung donors.** **(A)** Healthy donor 7664. **(B)** IPF donor 1015 with 60% fibrosis. **(C)** IPF donor 1033 with 90% fibrosis. Scale bar represents 1 mm.

**Figure S4 Individual violin plots for all coincident regions of the healthy and IPF human lung sections.** The individual violin plots quantify the heterogeneity of interactive states for AXL-GAS6 and AXL-PROS1 in **(A)** Healthy donor 7664, **(B)** IPF donor 1015, and **(C)** IPF donor 1033 lung sections. The dotted line at 4% Ef represents the Förster Radius ($R_0$). Values below 4% are not representative of ligand-receptor interactions.

**Figure S5 Validation of AAV6.2 and AAV9 tropisms for epithelial cells in mouse lung.** Experimental setup to prove the transduction specificity of AAV6.2 and AAV9. AAV6.2-GFP/ AAV9-GFP were administered (i.t) for tracking purposes. 14 days after AAV application, mice were sacrificed for histological staining. **(B-C)** GFP-staining of mouse lung sections transduced with AAV6.2-GFP and AAV9-GFP. **(B)** AAV6.2 serotype had specificity for airway/bronchial epithelial cells (BEC) and alveolar cells (AT-2 and AT-1 cells). **(C)** AAV9 serotype had specificity for alveolar cells. Brown color indicates GFP-positive staining. Box with dash line indicates areas that are zoomed in, while Grey arrow indicates BEC. Dot-like structure indicated by red arrow is AT-2 cell. The thin line-like structure indicated by blue arrow is AT-1 cell. Scale bar: 100 µm. Images are representative of n=4.

| Age (Years) | Sex | Diagnosis |
|---|---|---|
| 33 | Female | Non-diseased |
| 34 | Male | Non-diseased |
| 37 | Male | Non-diseased |
| 39 | Male | Non-diseased |
| 41 | Male | COPD |
| 44 | Female | COPD |
| 47 | Female | COPD |
| 57 | Female | COPD |
| 62 | Female | ILD |
| 69 | Male | IPF |
| 74 | Male | IPF |
| 76 | Male | ILD |
| 77 | Female | IPF |
| 83 | Male | ILD |
| 83 | Male | ILD |
| 84 | Female | ILD |

**Table S1. Demographic information of human BALF donors.** Donors included both male and female individuals. Non-diseased donors (purchased from Tissue Solutions, Ltd. UK) ranged in age from 33 – 39 years old. COPD donors (purchased from Tissue Solutions, Ltd. UK) ranged in age from 41 – 57 years old. ILD/ IPF donors (received from Wangen hospital, Germany) ranged in age from 62 – 84 years old.

#### Supplementary Methods

##### ELISA

ELISA were performed according to manufacturer's protocol. GAS6 and PROS1 was measured in human BALF or SAEC supernatant using the human GAS6 DuoSet ELISA kit (R&D Systems, #DY885B) and the human PROS1 ELISA Kit (Aviva Systems Biology, #OKBB010207). SP-D was measured in plasma using the mouse SP-D Quantikine ELISA kit (R&D Systems, #MSFPD0)

##### Western blot

AXL knockout SAEC was validated by Western blotting. Cells were lysed in RIPA buffer (Sigma Aldrich, #R0278-500ML) supplemented with protease- (Roche, #05892970001) and phosphatase (Roche, #04906837001) inhibitor. Protein concentrations were determined by BCA assay (ThermoFisher, #23225). Equal amounts of protein were denatured, resolved on 4-12% Bis-Tris gels (ThermoFisher, #NW04125BOX), and transferred to nitrocellulose membranes. Membranes were blocked and incubated overnight with antibodies against AXL 1:200 (R&D, #AF154) and ß-actin 1:1000 (Cell Signaling, #8457), followed by fluorescent secondary antibodies (donkey anti goat IR dye® 680 LT 1:5000 (LI-COR, #926-68021) and donkey anti rabbit IR dye® 800 CW 1:5000 (LI-COR, #926-32213). Signals were detected using an Odyssey Imager and quantified with LI-COR software.

##### Tissue dissociation

The mice were euthanized using pentobarbital (Narcoren, Boehringer-Ingelheim) overdose and the lungs were perfused with 10 ml RPMI (ThermoFisher, #11835030) through the right ventricle of the heart. The lungs and tracheas were harvested and placed into ice-cold RPMI. The tissues were minced with scissors into 1 mm size in 24 well plate and digested with 6.8 U/ml elastase (Merck KGAa, #324682-1000U), 0.3 Wünsch/ml liberase DL (Roche, #05466202001), and 0.1 mg/ml DNAse I (Roche, #10104159001) in 2.4 ml RPMI supplemented with 1% FBS (ThermoFisher #16140071) and 0.1 mM EDTA (ThermoFisher, #15575020). The tissues with the digestion mix were incubated for 90 minutes on a thermoblock at 37°C with 300 rpm agitation. The tissues with the enzyme solution were then pipetted with 2.5 ml pipette into a 100 µm strainer and pushed with the rubber end of a syringe plunger to obtain a single cell suspension. The strainer was washed with 10 ml RPMI supplemented with 1% FBS, 0.1 mM EDTA, and 0.1 mg/ ml DNAse I. The cells were pelleted by centrifugation at 300g for 5 minutes. If there were still red blood cells (RBC) visible, RBC lysis was performed for 2 minutes at RT using red blood cell lysing buffer Hybri-Max™ (Sigma, #R7757-100ML). After the washing step, the cell suspension was resuspended in PBS

(ThermoFisher #14190094) supplemented with 1% FBS. This protocol was adapted and modified from Donati, Y. *et al.* (2020)[31] .

##### Flow Cytometry

Lung and trachea cell suspension was incubated with fluorochrome conjugated antibody for extracellular staining with CD45 PerCp (BD Biosciences, #561047), CD31 PE-Cy7 (BD Biosciences, #561410), CD326 (EpCAM) BB515 (BD Biosciences, #565425), Pdpn PE (BD Biosciences, #566390), I-A/I-E (MHC-II) BV786 (BD Biosciences, #742894), and CD24 AF700 (BD Biosciences, #564237). The cells were fixed with BD Cytofix/Cytoperm™ and then subjected to intracellular staining with Ki67 AF647 (BD Biosciences, #561126). The antibody mix was supplemented with CD16/CD32 (Fc-block) (BD Biosciences, #553141). Fluorescence minus one (FMO) controls were included in all measurements. Acquisition was performed on BD LSR Fortessa X-20. FlowJo was used for data analysis.

##### Magnetic activated cell sorting (MACS)

Lung and trachea cell suspension was subjected to MACS according to the manufacturer's protocol (Miltenyi Biotec). Briefly, the cells were labelled with CD45 microbeads (Miltenyi Biotec, #130-052-301) to separate $CD45^+$ and $CD45^-$ cells. The $CD45^-$ fractions were subsequently labelled with EpCAM (CD326) (Miltenyi Biotec, #130-105-958) microbeads to enrich epithelial cells.

##### RNA Isolation for mRNA sequencing (mRNA-seq)

$EpCAM^+$ cells or SAEC were lysed in 350 µl RLT (Qiagen, #79216) supplemented with 1% 2-mercaptoethanol (Sigma Aldrich, #M6250). Lysate was pipetted into Qiaschredder (Qiagen, #79656) and centrifuged for 2 minutes at full speed. Phenol-Chloroform extraction was performed on EpCAM+ cell lysate, and the RNA was isolated using RNeasy micro kit according to the manufacturer's protocol (Qiagen, #74004)). While SAEC cell lysates were transferred into a MagMAX™ deep well plate and MagMAX™ mirVana™ Total RNA Isolation Kit (ThermoFisher, #A27828) was used alongside with the MagMAX™ Express-96 Deep Well Magnetic Particle Processor according to the manufacturer's protocol (ThermoFisher). Total RNA was quantitatively and qualitatively assessed using the fluorescence Broad Range Quant-iT RNA Assay Kit (ThermoFisher) and the Standard Sensitivity RNA Analysis DNF-471 Kit on a 96-channel Fragment Analyzer (Agilent), respectively. Total RNA samples had a RIN >7.5 and an input of 25ng (EpCAM+ cells) or 250ng (SAECs) was employed for bulk mRNA-seq library preparation with the NEBNext Ultra II Directional RNA Library Prep Kit for Illumina (#E7760), NEBNext Poly(A) mRNA Magnetic Isolation Module (#E7490) and NEBNext Multiplex Oligos for Illumina (#E7600) as per manufacturer's instructions (New England

Biolabs). Ampure XP beads (Beckman Coulter) for double-stranded cDNA purification were used instead of the recommended SPRIselect Beads. Libraries were amplified with 15 (EpCAM+ cells) or 13 (SAECs) PCR cycles and quantified with the High Sensitivity dsDNA Quanti-iT Assay Kit (ThermoFisher) on a Synergy HTX (BioTek). Libraries were assessed for size distribution and adapter dimer presence (<0.5%) by the High Sensitivity Small Fragment DNF-477 Kit on a 96-channel Fragment Analyzer (Agilent). Libraries were normalized on the MicroLab STAR (Hamilton), pooled and sequenced on a NovaSeq 6000 (Illumina) with dual index, paired-end reads (Read Parameter: Rd1:101, Rd2:10, Rd3:10, Rd4:101) with an average sequencing depth of >25 million Pass-Filter reads per sample.

**mRNA-seq analysis**

Briefly, demultiplexing was performed using bcl2fastq v2.20.0.422 from Illumina (https://emea.support.illumina.com/downloads/bcl2fastq-conversion-software-v2-20.html). Sequencing reads from the RNA-seq experiment were processed with a pipeline building upon the implementation of the ENCODE "Long RNA-seq" pipeline[32] , filtered reads were mapped against the Mus musculus (mouse) genome mm10/GRCm38 or the Homo sapiens (human) genome hg38/GRCh38 (primary assembly, excluding alternate contigs), respectively using the STAR (v2.5.2b) aligner[33] allowing for soft clipping of adapter sequences. For quantification, transcript annotation files from Ensembl version 86 were used, which corresponds to GENCODE M11 for mouse and GENCODE 25 for human. Gene expression levels were quantified with the above annotations using RSEM (v1.3.0)[34] and featureCounts (v1.5.1)[35]. Quality controls were implemented using FastQC (v0.11.5) [Andrews, S. (2010): http://www.bioinformatics.babraham.ac.uk/projects/fastqc/], picardmetrics (v0.2.4) [Slowikowski K. (2016): https://github.com/slowkow/picardmetrics] and dupRadar (v1.0.0)[36] at the respective steps. Finally, differential expression analysis was performed on the mapped counts derived from featureCount[36] using limma/voom[37]. An absolute log2 fold change cut-off of 1 and a false discovery rate (FDR) of <0.01 were applied.

**Ingenuity Pathway Analysis (IPA)**

Pathway analysis was performed with Qiagen Ingenuity Pathway Analysis IPA[38] [https://www.qiagenbioinformatics.com/products/ingenuity-pathway-analysis]. Significantly regulated genes from the pre-processing for each day and virus combination were used for both the canonical pathway analysis, excluding metabolic pathways, and the upstream analysis, focusing on the molecule type of transcriptional regulators with a predicted activation state. The results, for the analysis of the individual viruses over time on the pathway and upstream level, were filtered for both increasing and decreasing z-score values per pathway over time and sorted by the last or first time point, respectively. For the comparison between AAV6.2 and AAV9 at each timepoint, the top pathways and upstream regulators were selected based on sorting the combined z-scores, but including values for the selected categories, if they existed outside the top selection for one of the viruses.

**Cigarette smoke exposure in mice**

Mice were exposed to cigarette smoke over 28 days, starting with daily exposure for 7 days in the first week, followed by 5 days per week thereafter. Exposure was performed in a custom whole-body chamber equipped with an automated cigarette lighter, smoke generator, and heating pad. Mice received 2 cigarettes (without filters, tar 10 mg, nicotine 1.0 mg, carbon monoxide 6 mg) (Roth Händle) on day 1-2, 3 cigarettes on day 3-4, 4 cigarettes on day 5-6 for adaptation phase, and 5 cigarettes per day for the remainder of the study. Each cigarette last for 15 min, followed by 8-minutes fresh air (15 $L \cdot min^{-1}$), and an additional 23-minutes break after 2 cigarettes.
